# Mechanical Netrin-1-DCC Coupling Mediates Netrin-1–induced Axonal Haptotaxis

**DOI:** 10.1101/2023.12.03.569828

**Authors:** Zhen Qiu, Takunori Minegishi, Daichi Aoki, Kouki Abe, Kentarou Baba, Naoyuki Inagaki

## Abstract

The growth cone, a motile structure located at the tip of growing axon, senses extracellular guidance cues and translates them into directional forces that drive axon outgrowth and guidance. Axon guidance directed by chemical cues on the extracellular adhesive substrate is termed haptotaxis. Recent studies reported that netrin-1 on the substrate functions as a haptotactic axon guidance cue. However, the mechanism mediating netrin-1–induced axonal haptotaxis remains unclear. Here, we demonstrate that substrate-bound netrin-1 induces axonal haptotaxis by facilitating physical interactions between the netrin-1 receptor, DCC, and the adhesive substrates. DCC serves as an adhesion receptor for netrin-1. The clutch molecule shootin1a interacted with DCC, linking it to actin filament retrograde flow at the growth cone. Speckle imaging analyses showed that DCC underwent either grip (stop) and retrograde slip on the adhesive substrate. The grip state was more prevalent on netrin-1-coated substrate compared to the control substrate polylysine, thereby transmitting larger traction force on the netrin-1-coated substrate. Furthermore, disruption of the linkage between actin flow and DCC by shootin1 knockout impaired netrin-1-induced axonal haptotaxis. These findings indicate that the directional force for netrin-1–induced haptotaxis is exerted on the substrates through the mechanical coupling between netrin-1 and DCC which occurs asymmetrically under the growth cone.

## 1. Introduction

To establish functional neural networks, axons must navigate to appropriate synaptic partners. The growing tip of axons bears a motile structure the growth cone, which plays critical roles in axon outgrowth and guidance (Dent and Gertler, 2003; Lowery and Van Vactor, 2009; Vitriol and Zheng, 2012). Regarding the force to drive axon outgrowth and guidance, actin filaments (F-actins) polymerize at the leading edge of growth cones and disassemble proximally. This, together with the contractile activity of myosin II, drives retrograde flow of F-actin (Forscher and Smith, 1988; Katoh et al., 1999; Medeiros et al., 2006). Mechanical coupling between the F-actin flow and the adhesive substrate, mediated by “clutch” molecules and cell adhesion molecules, transmits the force of F-actin flow to the substrate as traction force for growth cone migration (Mitchison and Kirschner, 1988; Suter and Forscher, 2000; Shimada et al., 2008; Giannone et al., 2009). Concurrently, the actin-adhesion coupling slows down the F-actin flow (Suter et al., 1998; Toriyama et al., 2013), allowing actin polymerization to protrude the leading-edge membrane (Mogilner and Oster, 1996).

Axon guidance is regulated by chemical cues in the extracellular environment, including soluble chemicals (termed chemotaxis) and substrate-bound chemicals (haptotaxis) (Lowery and Van Vactor, 2009; Abe et al., 2018). Netrin-1 is one of the most well-characterized molecules for axon guidance (Ishii et al., 1992; Serafini et al., 1994; Lai Wing Sun et al., 2011; Boyer and Gupton, 2018); intensive research over the past decades has elucidated the molecular mechanics for netrin-1–induced axonal chemotaxis. Extracellular netrin-1 gradients, formed through diffusion, induce attraction of growth cone in vitro (Kennedy et al., 1994; Hong et al., 1999; Fothergill et al., 2014; Taylor et al., 2015; Baba et al., 2018). Stimulation of the netrin-1 receptor deleted in colorectal cancer (DCC) activates Cdc42 and Rac1 and their downstream kinase Pak1 in the growth cone (Li et al., 2002; Shekarabi and Kennedy, 2002; Shekarabi et al., 2005; Briançon-Marjollet et al., 2008; Demarco et al., 2012). Shootin1a functions as a clutch molecule that links F-actin retrograde flow and adhesive substrates by interacting with the actin-binding protein cortactin and the cell adhesion molecule L1 at the growth cone (Shimada et al., 2008; Kubo et al., 2015; Baba et al., 2018). Pak1, activated through the netrin-1–induced signaling, phosphorylates shootin1a (Toriyama et al., 2013). This phosphorylation enhances its binding affinity for cortactin and L1, thereby facilitating the traction force for netrin-1–induced axon outgrowth and chemoattraction (Kubo et al., 2015; Baba et al., 2018).

Recent studies have highlighted netrin-1, present on the adhesive substrate, as a haptotactic axon guidance cue (Mai et al., 2009; Moore et al., 2009; Moore et al., 2012; Dominici et al., 2017; Varadarajan et al., 2017). However, the molecular mechanics of netrin-1–induced haptotaxis remains unclear (Wu et al., 2019). Here, we report that DCC and shootin1a also mediate netrin-1–induced axonal haptotaxis, through a process that does not rely on cell signaling. Shootin1a interacted with DCC, linking it to F-actin retrograde flow. Speckle imaging analyses showed that DCC underwent either grip (stop) or retrograde slip (Abe et al., 2018) on the adhesive substrate. The grip state was more prevalent on netrin-1-coated substrate compared to the control substrate polylysine, leading to increased traction force transmission on the netrin-1-coated substrate. Furthermore, shootin1 knockout (KO) disrupted netrin-1–induced axonal haptotaxis. These findings indicate that the directional force for netrin-1–induced haptotaxis is exerted on the substrates through the asymmetric mechanical coupling between netrin-1 and DCC within the growth cone.

## 2. Materials and Methods

### 2.1. Animals

All relevant aspects of the experimental procedures were approved by the Institutional Animal Care and Use Committee of Nara Institute of Science and Technology. Shootin1 KO mice were generated as described previously (Baba et al., 2018). E16.5 shootin1 KO embryos were obtained by crossing male and female shootin1 heterozygous C57BL/6 mice; the offspring genotypes were confirmed by PCR using the following primers: Genotyping F1 (5′-CAGACTGCTACCCACTACCCCCTAC-3′) and Genotyping R1 (5′-CCTAGAGCTGGACAGCGGATCTGAG-3′) for the wild-type (WT) allele. Genotyping F2 (5′-CCCAGAAAGCGAAGGAACAAAGCTG-3′), Genotyping R2 (5′-ACCTTGCTCCTTCAAGCTGGTGATG-3′) for the KO allele. Embryonic stages were calculated from noon of the vaginal plug day, defined as embryonic day 0.5 (E0.5). Chimeric mice were crossed with C57BL/6 mice for at least nine generations before analysis. All C57BL/6 mice were bred under standard conditions (12 h/12 h light/dark cycle, access to dry food and water).

### 2.2. Cell culture and transfection

Hippocampal neurons were prepared form E16.5 WT and shootin1 KO mice as described (Minegishi et al., 2021). They were cultured on glass coverslips (Matsunami) or glass bottom dishes (Matsunami), which we coated with 100 μL/mL polylysine (poly-D-lysine) (Sigma, catalog number P6407-5MG) alone or sequentially with 100 μL/mL polylysine and 400 ng/mL netrin-1 (R&D systems, catalog number 6419-N1-025/CF), and cultured in Neurobasal Medium (Thermo Fisher Scientific) containing B-27 supplement (Thermo Fisher Scientific) and 1 mM glutamine. Micro-scale patterns of netrin-1 on polylysine-coated coverslips were prepared as described (Abe et al., 2018), using PBS solution containing 1M sucrose, 20 μg/ mL Texas Red-conjugated BSA (Thermo Fisher Scientific), and 400 ng/mL netrin-1 instead of 25 μg/ mL laminin. Briefly, a polydimethylsiloxane (PDMS) stamp was produced using a silicon substrate with small hexagonal patterns (Figure S1A). Then micro-scale patterns of netrin-1 were formed on polylysine-coated coverslips by a wet-transfer process using the PDMS stamp. All experiments, except for force measurements, were carried out on glass surfaces. Neurons were transfected with plasmid DNA using Nucleofector (Lonza) before plating as described (Minegishi et al., 2021). HEK293T cells (ATCC, RRID: CVCL_0063) were cultured in Dulbecco’s modified Eagle’s medium (Sigma) containing 10% fetal bovine serum (FBS) (Thermo Fisher Scientific) as described previously (Baba et al., 2018), and transfected with vectors using Polyethyleneimine MAX (Polysciences) according to the manufacturer’s protocol.

### 2.3. DNA constructs

To generate pCMV-FLAG-shootin1a, the cDNA of human shootin1a was amplified by PCR with the primers Shootin1a F (5′-CCGCTCGAGATGAACAGCTCGGACGAGGAGAAG-3′) and shootin1a R (5′-CCGCTCGAGTTACTGGGAGGCCAGGATTCCCTTCAG-3′), and then subcloned into pCMV-FLAG vector (Toriyama et al., 2006). To generate pFC14K-DCC-Halotag, the cDNA of human DCC was amplified by PCR with the primers DCC F (5′-GAGGATCCATGGAGAATAGTCTTAGATG-3′), and DCC R (5′-CTGGATCCCTAAAAGGCTGAGCCTGTGATGG-3′), and then subcloned into pFC14K-Halotag CMV Flexi vector (Promega). To generate pCMV-EGFP-DCC ICD (intracellular domain), the cDNA of human DCC was amplified by PCR with the primers DCC ICD F (5′-GGTGAACTTCAAGATCCGCCACAA-3′), and DCC ICD R (5′-TGCAATAAACAAGTTAACAACAACAATTG-3′), and then subcloned into pCMV-EGFP vector (Agilent Technology). The preparation of pFN21A-HaloTag-actin has been described previously (Minegishi et al., 2018).

### 2.4. Immunocytochemistry and microscopy

Cultured neurons were fixed with 3.7% formaldehyde in Krebs buffer (118 mM NaCl, 4.7 mM KCl,1.2 mM KH_2_PO_4_, 1.2 mM MgSO_4_, 4.2 mM NaHCO_3_, 2 mM CaCl_2_, 10 mM glucose, 400 mM sucrose, 10 mM HEPES pH 7.0) for 20 min on ice and subsequently for 10 min at room temperature, followed by treatment for 15 min with 0.05% Triton X-100 in phosphate-buffered saline (PBS) on ice and 10% FBS in PBS for 1 h at room temperature. They were then incubated with primary antibody diluted in PBS containing 10% FBS overnight at 4°C. The following primary antibodies were used: rabbit anti-shootin1a (1:2000) (Baba et al., 2018) goat anti-DCC (1:1000) (Santa Cruz Biotechnology, catalog number sc-515834), and mouse anti-Tuj1 (anti-β Ⅲ-Tubulin) (1:1000) (Bio Legend, catalog number 801202) antibodies. Neurons were washed with PBS, and then incubated with secondary antibody diluted in PBS 1 h at room temperature. The following secondary antibodies were used: Alexa Fluor 488 conjugated donkey anti-goat (1:1000) (Thermo Fisher Scientific, catalog number AB_2534102), Alexa Fluor 594 conjugated donkey anti-rabbit (1:1000) (Jackson ImmunoResearch Laboratories, catalog number AB_2340621), and Alexa Fluor 488 conjugated goat anti-mouse (1:1000) (Thermo Fisher Scientific, catalog number AB_2534088) antibodies. After washing with PBS, immunostained cells were mounted with 50% (v/v) glycerol (Nacalai Tesque) in PBS. Fluorescence images were acquired using either a fluorescence microscope (BZ-X710, Keyence) equipped with 20 × 0.75 NA (Nikon) and imaging software (BZ-X viewer, Keyence) or a total internal reflection fluorescence (TIRF) microscope (IX81, Olympus) equipped with an EM-CCD camera (Ixon3, Andor), a UAPON 100 × 1.49 NA (Olympus) and imaging software (MetaMorph, Molecular Devices).

### 2.5. Immunoprecipitation and immunoblotting

Lysates of HEK293T cells were prepared as described (Baba et al., 2018). Immunoprecipitation and immunoblotting were performed as described previously (Shimada et al., 2008; Baba et al., 2018). The following primary antibodies were used in immunoblotting: rabbit anti-GFP (1:2000) (MBL, catalog number AB_591816) and mouse anti-FLAG (DDDDK) tag (1:4000) (MBL, catalog number AB_591224). The following secondary antibodies were used in immunoblotting: HRP-conjugated donkey anti-rabbit IgG (1:2000) (GE Healthcare, catalog number AB_772206), HRP-conjugated goat anti-mouse IgG (1:5000) (Thermo Fisher Scientific, catalog number AB_2534088).

### 2.6. Fluorescent speckle imaging and grip and slip analysis

Fluorescent speckle imaging and speckle tracking analysis of HaloTag-actin were performed as described previously (Minegishi et al., 2021). Fluorescent speckle imaging and speckle tracking analysis of DCC-HaloTag was performed as described (Abe et al., 2018), using neurons transfected with pFC14K-DCC-Halotag. Neurons were treated with HaloTag TMR ligand (50 nM) (Promega, catalog number G8251) in culture medium and incubated for 1 h at 37℃. The ligand was then washed out with PBS, and the cells were incubated with L15 medium (Thermo Fisher Scientific) including B27 supplement (Thermo Fisher Scientific) and 1 mM glutamine for 30 min at 37℃. The fluorescent speckles of DCC-HaloTag were observed using a TIRF microscope (IX81, Olympus) equipped with an EM-CCD camera (Ixon3, Andor), a complementary metal-oxide-semiconductor (CMOS) camera (ORCA Flash4.0LT; Hamamatsu), a UAPOM 100x 1.49 NA (Olympus), and imaging software (MetaMorph, Molecular Devices). Fluorescence images were acquired every 5 sec for 295 sec.

We use the terms “grip” and “slip” to describe the movement of DCC-HaloTag. “Grip vs. slip” was used by Jurado *et al*. to describe the degree of interaction between the cell adhesion and adhesive substrate in fish keratocytes (Jurado et al., 2005). Subsequently, Aratyn-Schau and Gardel used “frictional slip” to describe the experimentally observed retrograde movement of focal adhesions (including integrin) of human osteosarcoma cells on adhesive substrates (Aratyn-Schaus and Gardel, 2010). Based on their concept and terms, we previously called L1 molecules that undergo retrograde movement on adhesive substrates as in “slip” phase and those immobilized on the substrates as in “grip” phase (Abe et al., 2018; Abe et al., 2021); we follow this terminology here. For the grip and slip analysis, DCC puncta that were visible for at least 10 s (two intervals) were analyzed, and immobile signals were defined as DCC in grip phase while those that flowed retrogradely were defined as DCC in slip phase.

### 2.7. Traction force microscopy

Traction force microscopy was performed as described previously (Minegishi et al., 2021). Neurons were transfected with pEGFP-C1 vectors and cultured on polyacrylamide gels with embedded 200-nm fluorescent beads (Thermo Fisher Scientific) for 2 days. Time-lapse imaging of fluorescent beads and growth cone was performed at 37℃ using a confocal laser microscope (LSM710, Carl Zeiss) equipped with a C-Apochromat 63x/1.2 W Corr objective, and ZEN2009 software. Magnitude of the traction force under the growth cones were calculated as described previously (Minegishi et al., 2021).

### 2.8. Quantification and statistical analysis

Data handling and preparation of graphs was performed using Microsoft Excel 2016 (Microsoft). Statistical tests were performed using GraphPad Prism7 (GraphPad Software). For samples with more than 7 data points, the D′Agostino–Pearson normality test was used to determine whether the data followed a normal distribution. For cases in which the number of data points was between 3 and 7, the Shapiro‒Wilk test was used for the normality test. For cases in which the number of data points was between 3 and 7, the Shapiro‒Wilk test was used for the normality test. We also tested the equality of variation with the F test for two independent groups that followed normal distributions. Significance tests were performed using two-tailed unpaired Student′s *t* test to compare two independent groups that followed normal distributions with equal variation. All data are shown as the mean ± SEM. Statistical significance was defined as ***p < 0.01; **p < 0.02; *p < 0.05; ns, not significant. A *P* value less than 0.05 was considered to be statistically significant. For each experiment, the corresponding statistics information and number of samples are indicated in the figure legend. All experiments were performed at least three times and reliably reproduced. Investigators were blind to the experimental groups for each analysis, except biochemical analyses.

## 3. Results

### 3.1. Substrate-bound netrin-1 promotes traction force for axon outgrowth and haptotaxis

To examine the effect of substrate-bound netrin-1 on axonal extension, we first cultured mouse hippocampal neurons on glass coverslips coated with either polylysine plus netrin-1 or polylysine alone as a control substrate. As reported (Dominici et al., 2017; Varadarajan et al., 2017; Wu et al., 2019), substrate-bound netrin-1 promoted axon outgrowth (Figure 1A, B). In addition, neurons cultured on micro-scale patterns of netrin-1 on polylysine-coated coverslips extended axons preferentially on netrin-1 (Figure 1C), indicating that substrate-bound netrin-1 induces axon outgrowth and attraction (haptotaxis) of cultured hippocampal neurons. To analyze the mechanics for netrin-1–induced axon outgrowth and haptotaxis, we measured traction force produced by the axonal growth cone. Neurons were cultured on polyacrylamide gels coated with polylysine alone or polylysine plus netrin-1; 200-nm fluorescent beads were embedded in the gels. Traction force under the growth cones was analyzed by visualizing force-induced deformation of the elastic gel, which is reflected by displacement of the beads from their original positions (Minegishi et al., 2021). As reported (Chan and Odde, 2008; Toriyama et al., 2013), the beads under the growth cone moved dynamically (Figure 1D, Supplementary Video S1). The magnitude of the traction force generated on netrin-1 was 11.9 ± 1.1 pN/μm^2^ (n = 7 growth cones), which was significantly larger than on polylysine (6.1 ± 1.5 pN/μm^2^) (n = 7 growth cones) (Figure 1E), indicating that substrate-bound netrin-1 promotes traction force for axon outgrowth and haptotaxis.

**FIGURE 1.**
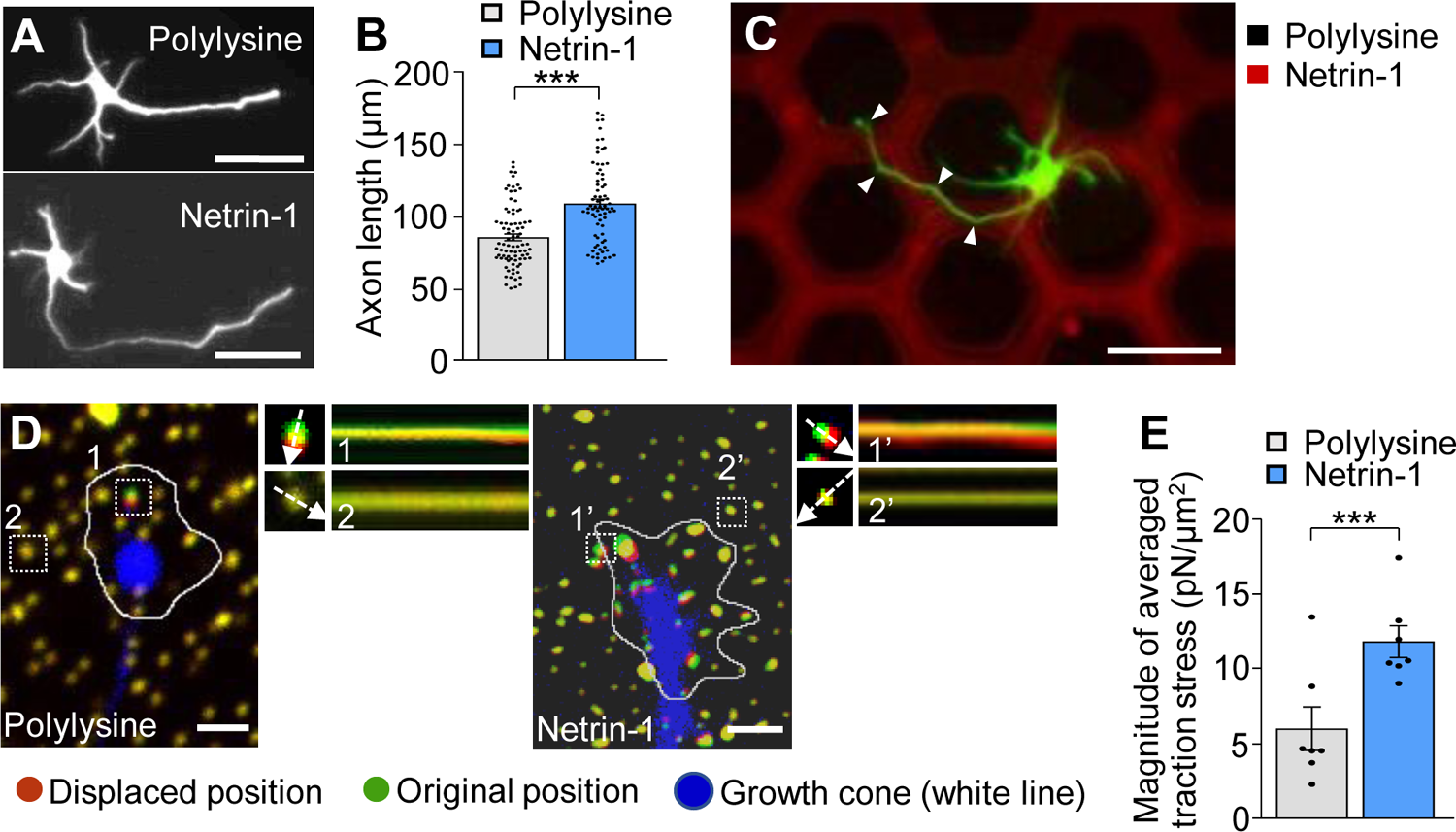
Substrate-bound netrin-1 promotes traction force for axon outgrowth and haptotaxis. (**A**) Fluorescence images of days *in vitro* (DIV) 2 hippocampal neurons cultured on glass coverslips coated with polylysine alone or coated sequentially with polylysine and netrin-1. Neurons were stained with a neuronal marker anti-Tuj1 antibody. (**B**) Quantification of axon length obtained by the analyses using neurons in (**A**) (polylysine, n = 85 cells; netrin-1, n = 75 cells). (**C**) A hippocampal neuron cultured on a micro-scale pattern of netrin-1 (red) and polylysine (black) for 2 days. Neurons were stained with anti-Tuj1 antibody (green). The axon turned toward netrin-1 when it reached the border between netrin-1 and polylysine (arrowheads). (**D**) Fluorescence images showing axonal growth cones of DIV 2 neurons expressing EGFP (blue) and cultured on polylysine-coated (left) or netrin-1-coated (right) polyacrylamide gel with embedded 200-nm fluorescent beads. The pictures show representative images from time-lapse series taken every 3 sec for 147 sec. White lines indicate the growth cone boundaries (see Supplementary Video S1). The kymographs along the axis of bead displacement (white dashed arrows) at the indicated areas show movement of beads recorded every 3 sec. The beads in areas 2 and 2’ are reference beads. (**E**) Quantification of the magnitude of the traction forces under axonal growth cones on polylysine-coated or netrin-1-coated polyacrylamide gels in (**D**) (Polylysine = 7 growth cones; Netrin-1 = 7 growth cones). Scale bars, 25 μm for (**A**); 50 μm for (**C**); 2 μm for (**D**). Data represent means ± SEM; ***p < 0.01.

### 3.2. Substrate-bound netrin-1 promotes F-actin-substrate coupling

Previous studies reported that the mechanical coupling between F-actin retrograde flow and the adhesive substrate promotes traction force (blue arrows, Figure 2A); concurrently, it reduces the F-actin flow speed (yellow arrows) (Suter and Forscher, 2000; Toriyama et al., 2013). To further examine the molecular mechanism of the netrin-1–induced force generation, we monitored F-actin retrograde flow within the axonal growth cone by fluorescent speckle imaging of HaloTag-actin (Minegishi et al., 2021). HaloTag-actin speckles moved retrogradely in the of growth cones (Figure 2B, Supplementary Video S2), depicting the F-actin movement (Figure 2A). The F-actin flow speed on polylysine was 4.5 ± 0.3 μm/min (mean ± SEM, n = 84 signals, 6 growth cones). On the other hand, netrin-l on substrates significantly reduced the flow speed (2.3 ± 0.1 μm/min, n = 92 signals, 6 growth cones) (Figure 2B, C; Supplementary Video S2). Together with the observed increase in traction force on netrin-1 (Figure 1E), these data indicate that netrin-1 on the adhesive substrate promotes the coupling between F-actin retrograde flow and the adhesive substrate at the growth cone.

**FIGURE 2.**
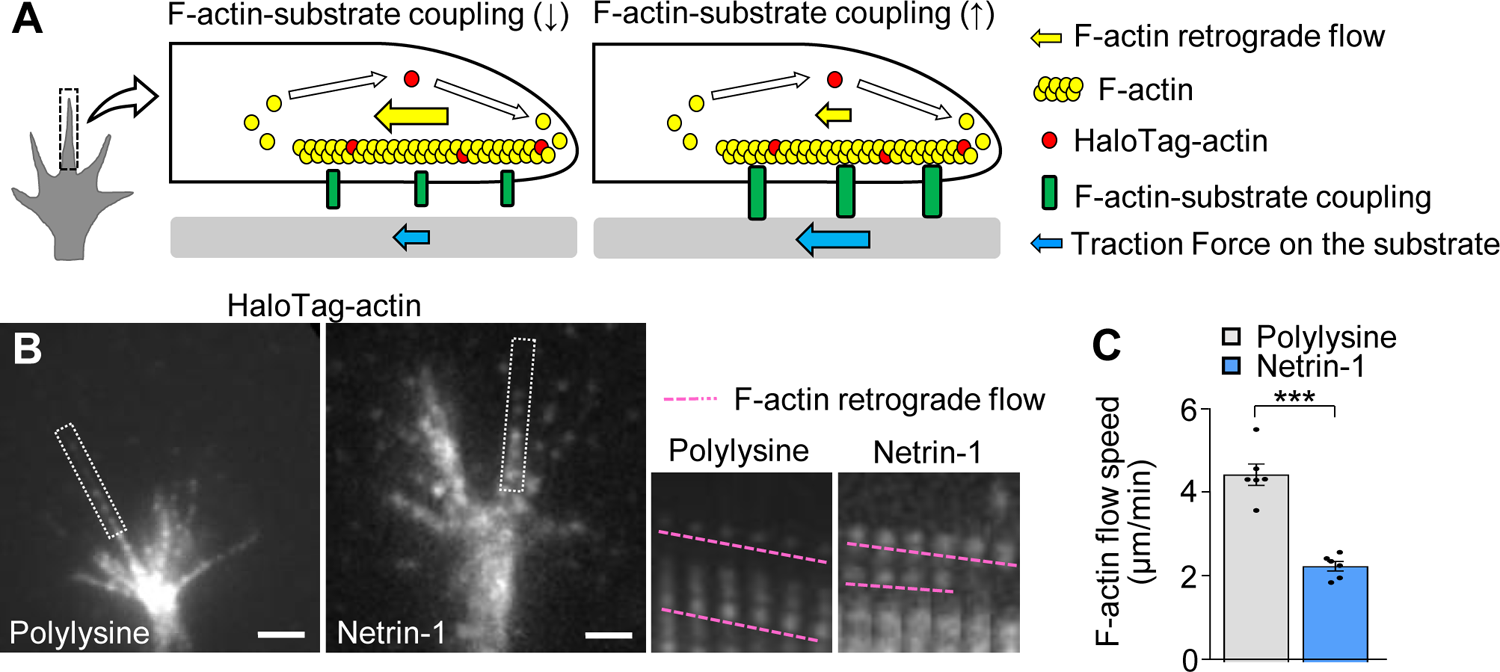
Substrate-bound netrin-1 promotes F-actin-substrate coupling. **(A)** A diagram explaining how the mechanical coupling between F-actin retrograde flow and the adhesive substrate generates traction force at the growth cone. Right: F-actin-substrate coupling (green) transmits the force of F-actin retrograde flow (yellow arrow) to the substrate as traction force (blue arrow) for growth cone migration. Left: a reduction in F-actin-substrate coupling decreases the force transmission (blue arrow), and increases the F-actin flow speed (yellow arrow) as the impediment to the flow by the coupling is diminished. (**B**) Fluorescent speckle images of HaloTag-actin in axonal growth cones cultured on polylysine (left) and netrin-1 (right); time-lapse montages of the fluorescent speckles of HaloTag-actin in filopodia on netrin-1 and on polylysine (boxed areas) at 5-sec intervals are shown to the right. F-actin flows are indicated by pink dashed lines (see Supplementary Video S2). (**C**) Quantification of F-actin flow speeds measured from the time-lapse montage analyses in (**B**) (polylysine, n = 84 signals, 6 growth cones; netrin-1, n = 92 signals, 6 growth cones). Scale bars, 2 μm for (**B**). Data represent means ± SEM; ***p < 0.01.

### 3.3. DCC and shootin1a couple F-actin flow with substrate-bound netrin-1

Next, we examined the molecular components linking F-actin retrograde flow to substrate-bound netrin-1. Previous studies reported that the clutch molecule shootin1a interacts with the intracellular domains of the immunoglobulin superfamily adhesion molecules, L1 and N-cadherin (Baba et al., 2018; Kastian et al., 2021). This led us to hypothesize that shootin1a interacts with another member of immunoglobulin superfamily, DCC (Lai Wing Sun et al., 2011). In addition, microbead tracking analyses reported that the mechanical interaction between the growth cone and substrate-bound netrin-1 exerts forces on the substrate in a DCC dependent manner (Moore et al., 2009). To assess the involvement of DCC and shootin1a in the linkage between F-actin flow and substrate-bound netrin-1, we performed co-immunoprecipitation assay, using HEK293T cells which express shootin1a and the intracellular domain of DCC (DCC-ICD). Shootin1a interacted with DCC-ICD (Figure 3A). In addition, shootin1a colocalized with DCC in the axonal growth cones of hippocampal neurons (Figure 3B), suggesting that shootin1a interacts with DCC-ICD at growth cones. Furthermore, disruption of shootin1a function by KO increased the F-actin flow speed at the growth cone on netrin-1-coated substrate (Figure 3C, D; Video S3). Concurrently, shootin1 KO reduced the magnitude of the traction force on the netrin-1-coated substrate (Figure 3E, F, Video S4). Together, we conclude that DCC and shootin1a couple F-actin flow with substrate-bound netrin-1 (Figure 3G).

**FIGURE 3.**
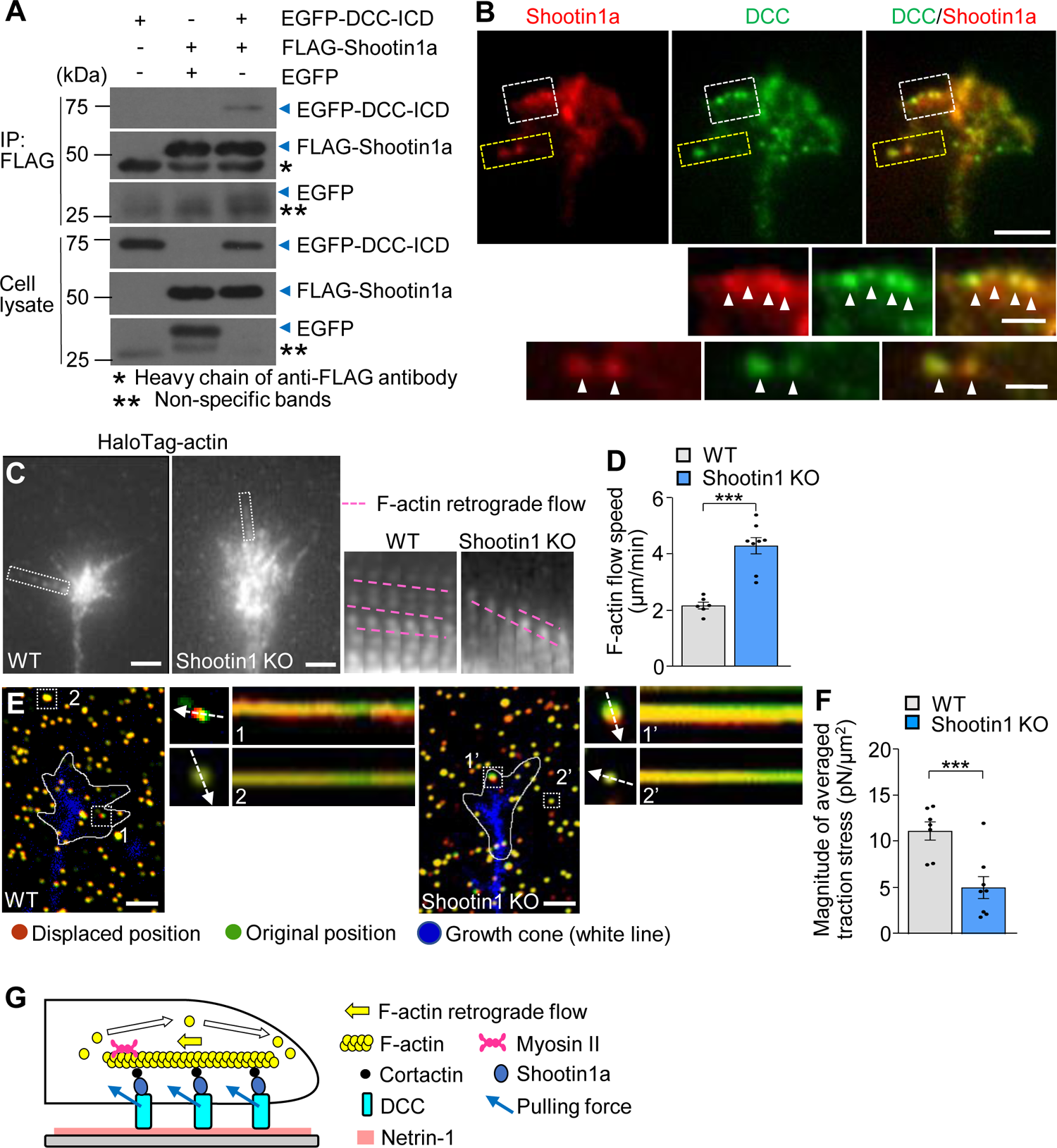
DCC and shootin1a couple F-actin flow with substrate-bound netrin-1. (**A**) Co-immunoprecipitation of DCC-ICD with shootin1a in HEK 293T cells. HEK293T cells were transfected with vectors to express FLAG-shootin1a and EGFP-DCC-ICD; cells were also co-transfected with a vector to express EGFP as a negative control. Cell lysates were prepared and incubated with anti-FLAG antibody for immunoprecipitation. The immunoprecipitants were immunoblotted with anti-FLAG or anti-GFP antibody. Whole immunoblot images are presented in Supplementary Figure S2. (**B**) Fluorescence images of axonal growth cone of a DIV 2 mouse hippocampal neuron labeled with anti-shootin1a (red) and anti-DCC (green) antibodies. Images below show enlarged views of the lamellipodium in the white square and the filopodium in the yellow square. Arrowheads indicate DCC colocalized with shootin1a. (**C**) Fluorescent speckle images of HaloTag-actin in axonal growth cones of WT (left) and shootin1 KO (right) neurons cultured on netrin-1. Time-lapse montages of HaloTag-actin in filopodia (boxed areas) at 5-sec intervals are shown to the right; pink dashed lines indicate the retrograde flow of speckles (see Supplementary Video S3). (**D**) Quantification of F-actin flow speeds measured from the time-lapse montage analyses in (**C**) (WT, n = 57 signals, 8 growth cones; shootin1 KO, n = 74 signals, 6 growth cones). (**E**) Fluorescence images showing axonal growth cones expressing EGFP (blue) of DIV 2 WT (left) and shootin1 KO (right) neurons cultured on netrin-1-coated polyacrylamide gel with embedded 200-nm fluorescent beads. The pictures show representative images from time-lapse series taken every 3 sec for 147 sec. White lines indicate the growth cone boundaries (see Supplementary Video S4). The kymographs along the axis of bead displacement (white dashed arrows) at the indicated areas show movement of beads recorded every 3 sec. The beads in areas 2 and 2’ are reference beads. (**F**) Quantification of the magnitude of the traction forces under axonal growth cones of WT and shootin1 KO neurons cultured on netrin-1-coated polyacrylamide gels in (**E**) (WT, n = 7 growth cones; shootin1 KO, n = 8 growth cones). **(G)** A diagram showing F-actin-substrate coupling in the axonal growth cone through cortactin, shootin1a, DCC and substrate-bound netrin-1. Shootin1a interacts with the actin binding protein cortactin (Kubo et al., 2015) and DCC (**A**). Scale bars: 5 μm for (**B**) upper figures; 1 μm for (**B**) enlarged figures; 2 μm for (**C**) and (**E**). Data represent means ± SEM; ***p < 0.01.

### 3.4. Netrin-1–induced axon outgrowth and haptotaxis require shootin1a-mediated actin-DCC coupling

To further examine the role of the actin-substrate linkage mediated by DCC and shootin1a, we analyzed axon outgrowth and haptotaxis. Disturbance of this linkage by shootin KO inhibited axon outgrowth on netrin-1-coated substrate (Figure 4A, B). As we have seen, hippocampal neurons cultured on micro-scale patterns of netrin-1 on polylysine-coated coverslips extended axons preferentially on netrin-1 as haptotaxis (Figure 4C); 79.6% ± 3.0% (n = 16 neurons) of the total lengths of axons were located on netrin-1-coated substrate (Figures S1B and 4D). On the other hand, shootin1 KO neurons extended axons which frequently crossed the borders between netrin-1 and polylysine (Figure 4C); they extended significantly lower rate (45.4% ± 3.4%, n = 14 neurons) of axons on netrin-1-coated substrate (Figure 4D). These data indicate that netrin-1–induced axon outgrowth and haptotaxis require actin-DCC coupling mediated by shootin1a (Figure 3G).

**FIGURE 4.**
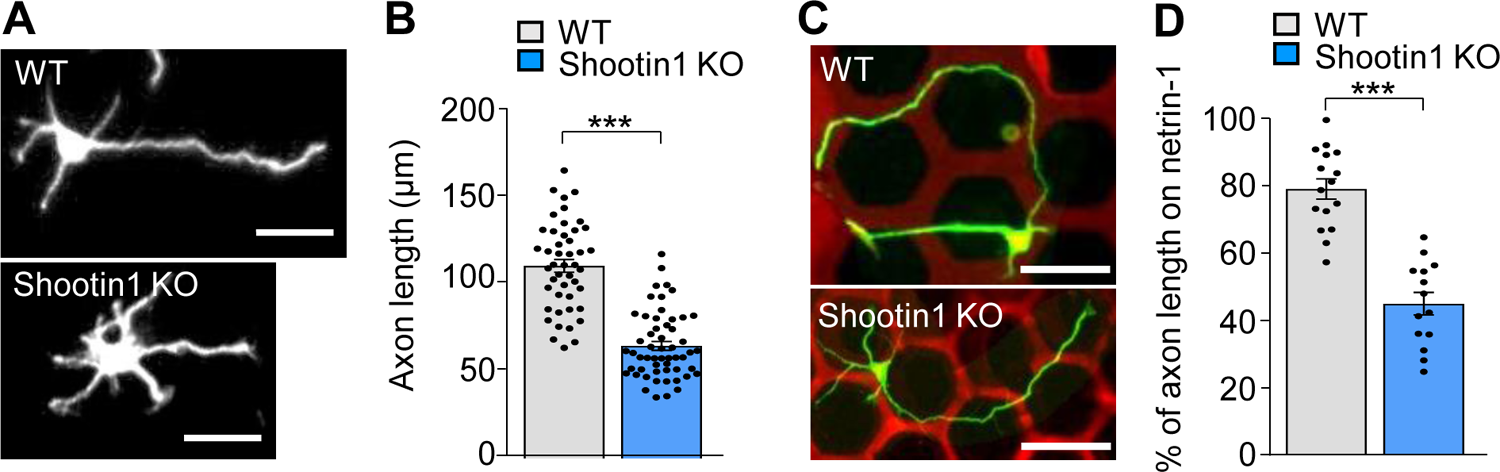
Netrin-1–induced axon outgrowth and haptotaxis require shootin1a-mediated actin-DCC coupling. (**A**) Fluorescence images of DIV 2 hippocampal neurons prepared from WT or shootin1 KO mouse and cultured on netrin-1. Neurons were stained with anti-Tuj1 antibody. (**B**) Quantification of axon length obtained by the analyses using neurons in (**A**) (WT, n = 46 cells; shootin1 KO, n = 55 cells). (**C**) Hippocampal neurons cultured on micro-scale patterns of netrin-1 (red) and polylysine (black) till DIV 3. Neurons were stained with anti-Tuj1 antibody (green). (**D**) Quantification of the percentage of axon length located on netrin-1 (see Supplementary Figure S1B) (WT, n = 30 cells; shootin1 KO, n = 14 cells). Scale bars, 25 μm for (**A**); 50 μm for (**C**). Data represent means ± SEM; ***p < 0.01.

### 3.5. DCC in growth cones undergoes differential grip and slip on the substrates

To analyze the mechanism by which substrate-bound netrin-1 promotes F-actin-substrate coupling, we monitored the movement of DCC, which serves as the link between intracellular shootin1a and netrin-1 on the substrate (Figure 3G). DCC-HaloTag expressed in hippocampal neurons was labeled with TMR ligand and observed using a TIRF microscope (Figure 5A). Two distinct types of DCC signals were observed: retrogradely flowing (blue dashed lines) and immobile (pink dashed lines, Figure 5A). We previously reported similar movements of L1 at the growth cone during laminin-induced axonal haptotaxis (Abe et al., 2018) and mechanosensing (Abe et al., 2021). At the growth cone, the force of F-actin retrograde flow is transmitted to L1 through cortactin and shootin1a, allowing to pull the bond between L1 and laminin. When the pulling force exceeds a threshold, the bond breaks and L1 at flows retrogradely on the substrate (Abe et al., 2018; Abe et al., 2021).

**FIGURE 5.**
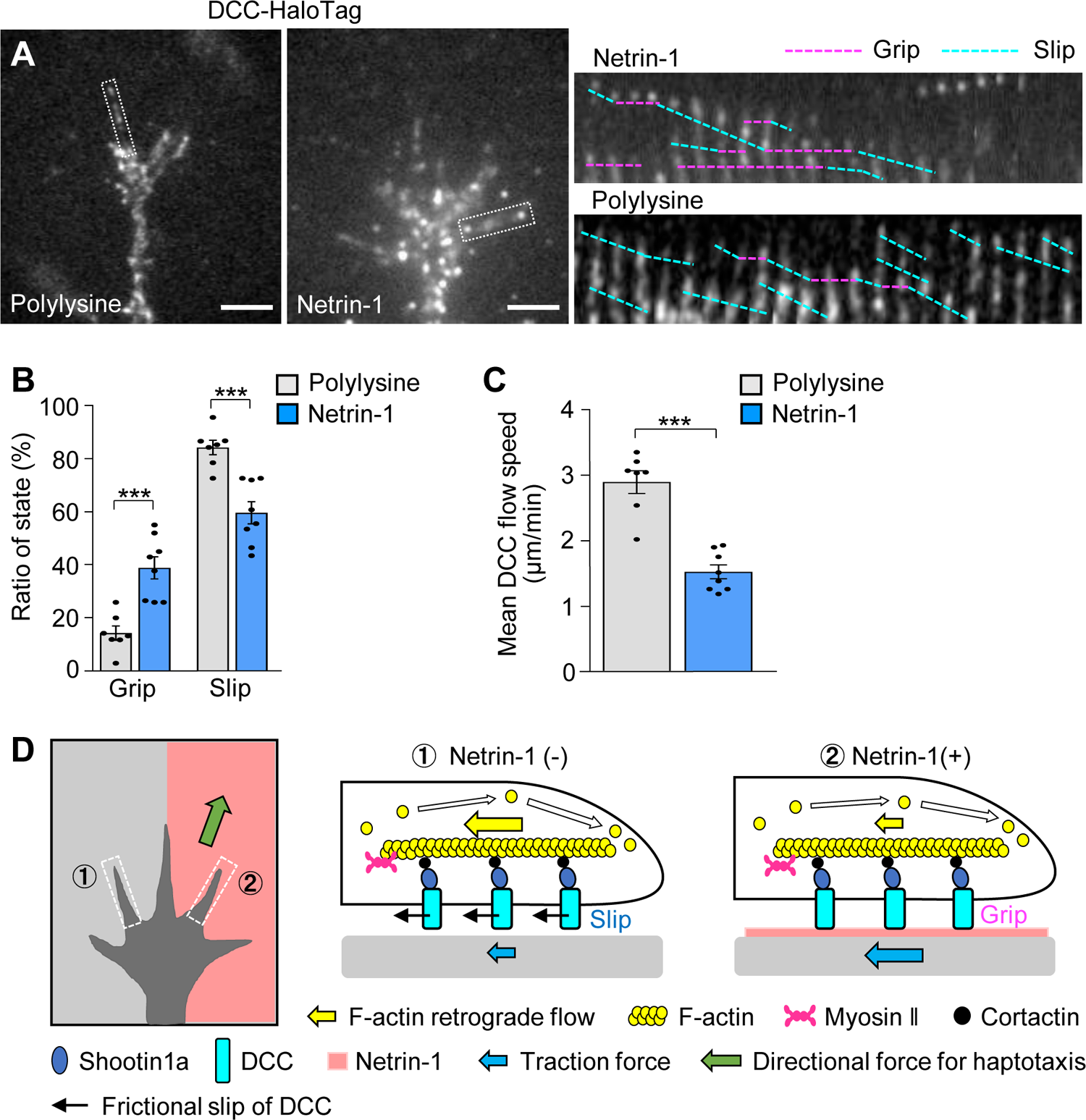
DCC in growth cones undergoes differential grip and slip on the substrates. (**A**) Fluorescent speckle images of DCC-HaloTag at growth cone membranes cultured on polylysine (left) and netrin-1 (right); time-lapse montages of DCC-HaloTag speckles in filopodia on netrin-1 and on polylysine (boxed areas) at 5-sec intervals are shown (grip and slip phases are indicated by dashed pink and blue lines, respectively) (see Supplementary Video S5). **(B** and **C)** Ratio of the grip and slip states (**B**) and retrograde flow speed (**C**) of DCC-HaloTag in filopodia measured from the time-lapse montage analyses in (**A**) (polylysine, n = 106 signals, 350 speckles, 7 growth cones; netrin-1, n = 114 signals, 548 speckles, 8 growth cones). (**D**) Grip and slip mechanism for netrin-1–induced axonal haptotaxis. The force of F-actin retrograde flow in the growth cone (yellow arrows) is transmitted to DCC through cortactin and shootin1a, allowing to pull the bond between DCC and adhesive substrates. DCC molecules undergo grip or frictional slip (black arrows) on the substrates. As the substrate presenting netrin-1 has higher affinity to DCC than the control substrate, a larger number of DCC molecules undergo grip on the netrin-1-presenting substrate compared to the control substrate, leading to more efficient force transmission on the netrin-1 side (blue arrows). This asymmetric grip and slip of DCC within the growth cone generate directional force for axonal haptotaxis toward netrin-1 (green arrow). Scale bars, 2 μm for (**A**). Data represent means ± SEM; ***p < 0.01.

We expect that similar force are transmitted to DCC, thereby pulling the bond between DCC and substrate-bound netrin-1 (blue arrows, Figure 3G). We refer to DCC that undergoes retrograde movement on the substrates as being in “slip” phase and that immobilized on the substrates as being in “grip” phase, following the terminology of L1 movements (see Materials and Methods) (Abe et al., 2018). On polylysine, 15% of DCC speckles exhibited the grip phase, while 85% displayed slip phase (n = 350 speckles, 7 growth cones). In contrast, 40% of the signals were in grip phase on netrin (n = 548 speckles, 8 growth cones) (Figure 5B). The grip phase percentage was significantly larger on netrin-1 than on polylysine, indicating that DCC binds to the netrin-1-coated substrate with higher affinity. Consistently, the mean flow speed of DCC on the netrin-1-coated substrate was 1.63 ± 0.11 μm/min (n = 114 signals, 8 growth cones), significantly slower than that on the polylysine-coated substrate (3.09 ± 0.18 μm/min, n = 106 signals, 7 growth cones) (Figure 5C). Together, these data suggest that netrin-1 promotes F-actin-substrate coupling at the interphase between DCC and the substrate, thereby generating traction forces for axon outgrowth and haptotaxis (Figure 5D).

## 4. Discussion

We have shown that DCC and shootin1a mechanically couple F-actin retrograde flow in the axonal growth cone and netrin-1 on the adhesive substrate. Substrate-bound netrin-1 increased the grip phase of DCC and promoted F-actin-substrate coupling, thereby increasing traction force produced by the growth cone. Furthermore, inhibition of the mechanical coupling by shootin1a KO disrupted netrin-1–induced axon outgrowth and haptotaxis. These data suggest a mechanism by which the growth cone generates a directional force for netrin-1–induced axonal haptotaxis (Figure 5D). The force of F-actin retrograde flow in the growth cone (yellow arrows) is transmitted to DCC through cortactin and shootin1a, allowing to pull the bond between DCC and the adhesive substrates. When the pulling force exceeds a threshold, the bond breaks and DCC molecule flows retrogradely on the substrate (black arrows). As the substrate presenting netrin-1 exhibits higher affinity to DCC than the control substrate, more DCC molecules grip the netrin-1-presenting substrate than the control, enabling more efficient force transmission on the netrin-1 side (blue arrows). This asymmetric gripping and slipping of DCC within the growth cone generate directional force for axonal haptotaxis toward netrin-1 (green arrow). The present axon guidance mechanism does not rely on cell signaling but depends on differences in the physical force between DCC and the adhesive substrates, with DCC serving as an adhesion receptor for netrin-1. This mechanism is consistent with the previous report that growth cones exerted a pulling force on netrin-1 in a DCC-dependent manner (Moore et al., 2009). Additionally, we recently reported a similar “grip and slip” mechanism for laminin-induced axonal haptotaxis, where difference in physical forces between the cell adhesion molecule L1 and the substrates mediated axon turning (Abe et al., 2018). Given its simplicity, we consider that this mechanism may contribute to various haptotactic axon guidance and cell migration. Versatility of this mechanism in haptotactic cell motility is an intriguing issue for further investigation.

DCC also serves as a cell signaling receptor for netrin-1 (Lai Wing Sun et al., 2011). Netrin-1 binding to DCC activates Cdc42, Rac1 and Pak1 (Li et al., 2002; Shekarabi and Kennedy, 2002; Shekarabi et al., 2005; Briançon-Marjollet et al., 2008; Demarco et al., 2012; Boyer and Gupton, 2018). Netrin-1–induced axonal chemoattraction is also mediated by various signaling molecules, including phosphatidylinositol-3 kinase, focal adhesion kinase, phospholipase Cγ, cAMP, Ca^2+^, ERK1/2, and Src. (Song and Poo, 2001; Gomez and Zheng, 2006; Lowery and Van Vactor, 2009; Lai Wing Sun et al., 2011; Moore et al., 2012; Gomez and Letourneau, 2014; Sutherland et al., 2014). In addition, shootin1a mediates netrin-1–induced axon guidance under the DCC signaling cascade; it is phosphorylated by Pak1 upon DCC stimulation and dephosphorylated by protein phosphatase-1 (Toriyama et al., 2013; Kastian et al., 2023). Phosphorylation of shootin1a enhances F-actin-substrate coupling and promotes the traction force for netrin-1–induced axon outgrowth and chemoattraction (Kubo et al., 2015; Baba et al., 2018). Netrin-1 gradients induce chemoattraction even without substrate-bound netrin-1, indicating that netrin-1 can function as a chemotactic guidance cue (Baba et al., 2018). The mechanism involving shootin1a phosphorylation is highly sensitive to extracellular gradients of netrin-1 (Baba et al., 2018), potentially involving desensitization and re-adaptation of the signaling pathway for long-distance axon guidance (Kastian et al., 2023). During navigation, growth cones may sense netrin-1 as an adhesion or cell signaling ligand for DCC, accommodating to different environments along their routes, to travel to their destinations.

## SUPPLEMENTARY MATERIAL

The Supplementary Material for this article can be found online at:

**SUPPLEMENTARY VIDEO S1** Time-lapse movies of axonal growth cones of hippocampal neurons expressing EGFP (blue) and cultured on polylysine-coated and netrin-1-coated polyacrylamide gels with embedded 200-nm fluorescent beads (see Figure 1D). Images were acquired every 3 sec for 147 sec using a confocal microscope.

**SUPPLEMENTARY VIDEO S2** Time-lapse movies of HaloTag-actin in axonal growth cones of hippocampal neurons cultured on polylysine-coated and netrin-1-coated glass-bottom dishes (see Figure 2B). Images were acquired every 5 sec for 295 sec using a fluorescence microscope.

**SUPPLEMENTARY VIDEO S3** Time-lapse movies of HaloTag-actin in axonal growth cones of a WT hippocampal neuron and a shootin1 KO hippocampal neuron cultured on netrin-1-coated glass-bottom dishes (see Figure 3C). Images were acquired every 5 sec for 295 sec using a fluorescence microscope.

**SUPPLEMENTARY VIDEO S4** Time-lapse movies of axonal growth cones of a WT hippocampal neuron and a shootin1 KO hippocampal neuron expressing EGFP (blue) and cultured on netrin-1-coated polyacrylamide gels with embedded 200-nm fluorescent beads (see Figure 3E). Images were acquired every 3 sec for 147 sec using a confocal microscope.

**SUPPLEMENTARY VIDEO S5** Time-lapse movies of DCC-HaloTag in axonal growth cones of hippocampal neurons cultured on polylysine-coated and netrin-1-coated glass-bottom dishes (see Figure 5A). Images were acquired every 5 sec for 245 sec using a TIRF microscope.

---

**SUPPLEMENTARY FIGURE S1.**
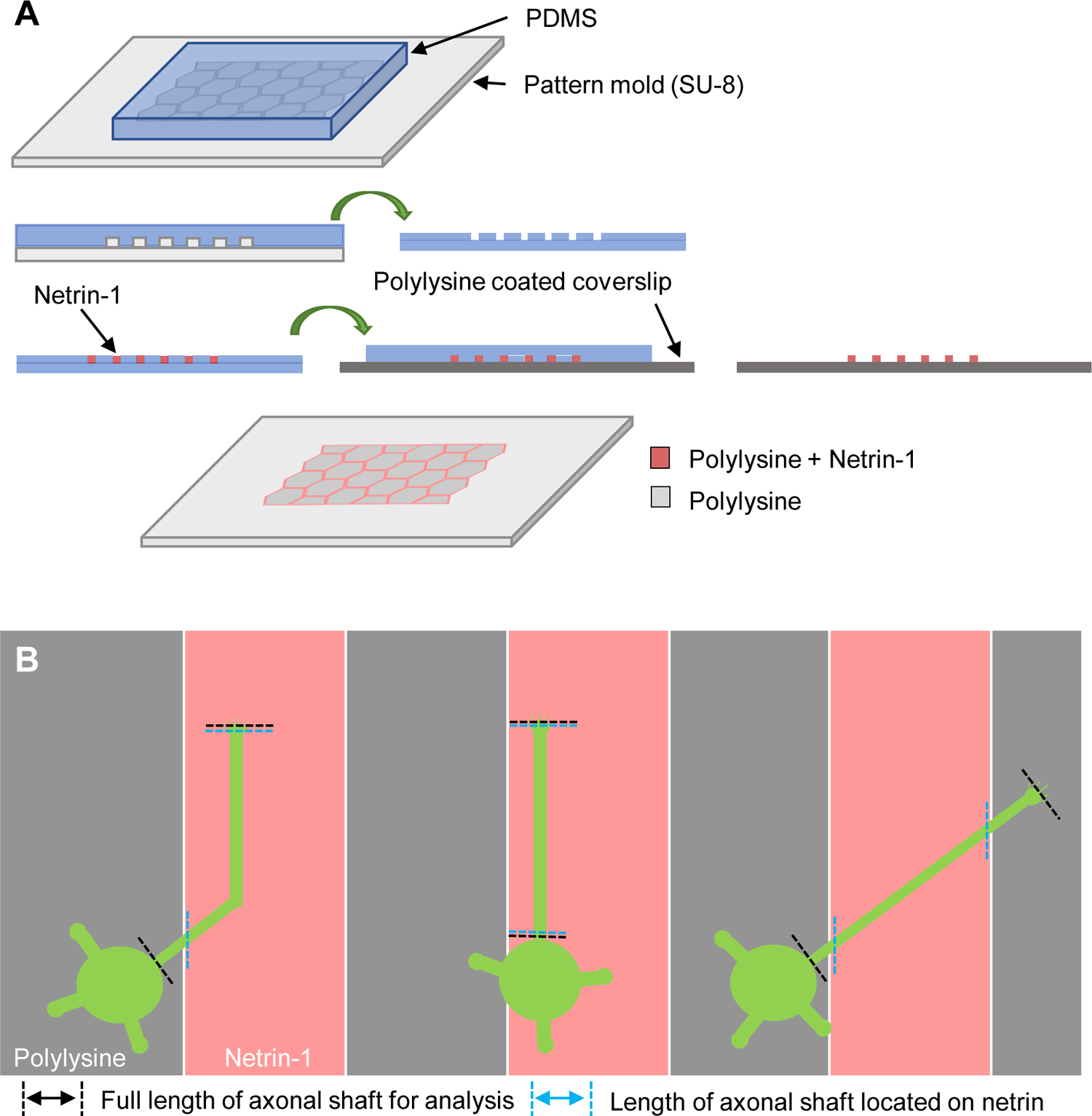
Analysis of netrin-1–induced axonal haptotaxis. (**A**) Preparation of micro-scale patterns of netrin-1 on polylysine-coated coverslips. For details, see Materials and Methods. (**B**) Analysis of axons located on netrin-1. To calculate % of axon length on netrin-1, the total lengths of axonal shafts located on netrin-1-coated areas (blue double arrow) were divided by the full lengths of the axonal shafts (black double arrow).

**SUPPLEMENTARY FIGURE S2.**
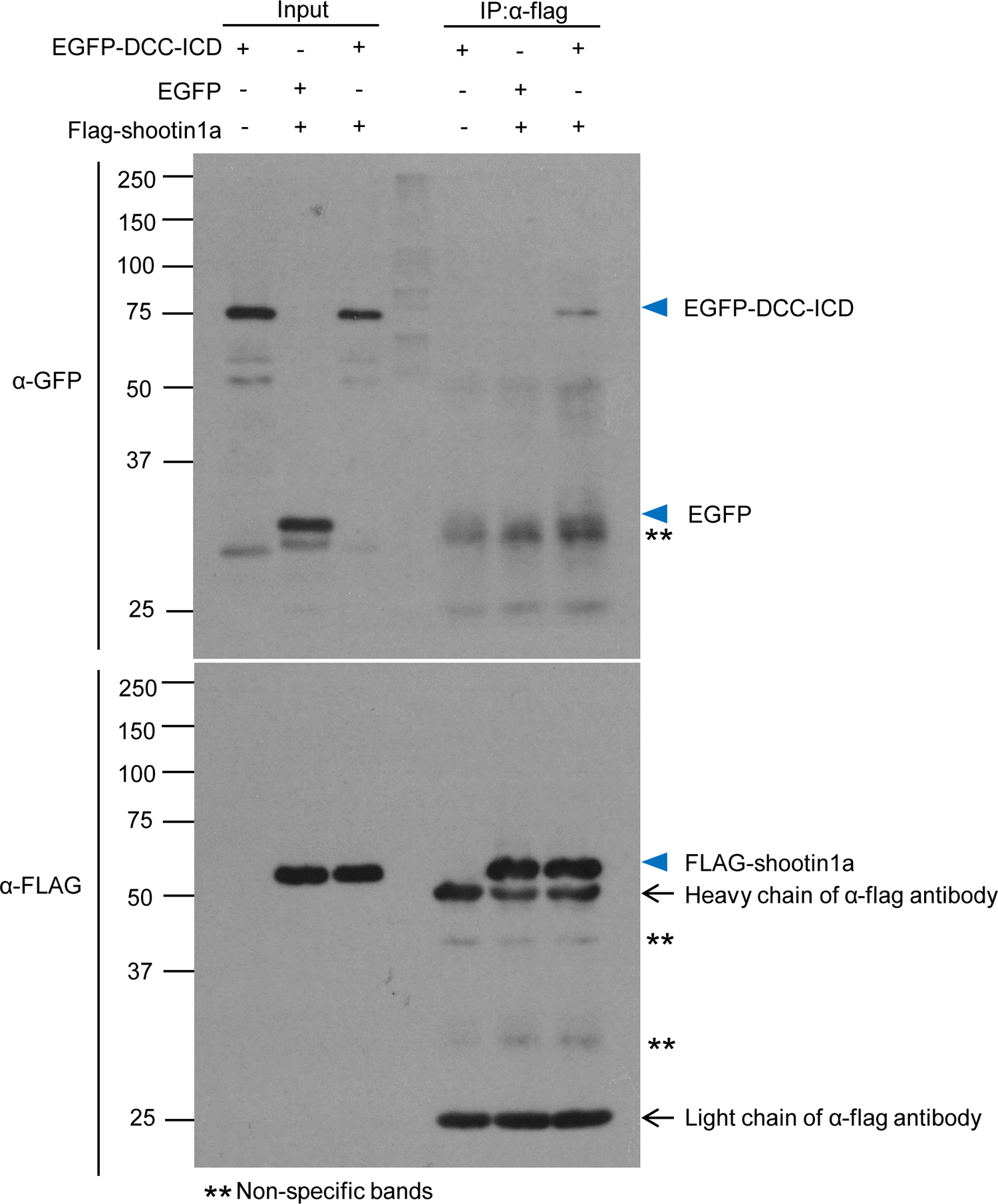
Whole immunoblot images in Figure 3A HEK293T cells were transfected with vectors to express FLAG-shootin1a and EGFP-DCC-ICD; cells were also co-transfected with a vector to express EGFP as a negative control. Cell lysates were prepared and incubated with anti-FLAG antibody for immunoprecipitation. The immunoprecipitants were immunoblotted with anti-FLAG or anti-GFP antibody.

## Data availability statement

The raw data supporting the conclusions of this article will be made available by the authors, without undue reservation.

## Author contributions

Z.Q., D.A. performed the experiments and data analysis. Z.Q., T.M., K.A., D.A., K.B., and N.I. designed the experiments. Z.Q., T.M. and N.I. wrote the manuscript. N.I. supervised all the projects. All authors discussed the results and commented on the manuscript.

## Acknowledgements

We thank Drs. Akira Kurisaki and Katsumoto Okamura (Nara Institute of Science and Technology) for discussions; Mieko Ueda and Kazumi Maekawa for technical support; Satoko Shimamura for kind encouragement; and Hongjia Chen, Kazuma Kami, and Singh Saranpal for supporting data analyses.

## References

Abe, K., Baba, K., Huang, L., Wei, K.T., Okano, K., Hosokawa, Y., et al. (2021). Mechanosensitive axon outgrowth mediated by L1-laminin clutch interface. Biophys. J. 120, 3566–3576. doi: 10.1016/j.bpj.2021.08.009.

Abe, K., Katsuno, H., Toriyama, M., Baba, K., Mori, T., Hakoshima, T., et al. (2018). Grip and slip of L1-CAM on adhesive substrates direct growth cone haptotaxis. Proc Natl Acad Sci U S A 115, 2764–2769. doi: 10.1073/pnas.1711667115.

Aratyn-Schaus, Y., and Gardel, M.L. (2010). Transient frictional slip between integrin and the ECM in focal adhesions under myosin II tension. Curr. Biol. 20, 1145–1153. doi: 10.1016/j.cub.2010.05.049.

Baba, K., Yoshida, W., Toriyama, M., Shimada, T., Manning, C.F., Saito, M., et al. (2018). Gradient-reading and mechano-effector machinery for netrin-1-induced axon guidance. Elife 7. doi: 10.7554/eLife.34593.

Boyer, N.P., and Gupton, S.L. (2018). Revisiting netrin-1: one who guides (axons). Front. Cell Neurosci. 12, 221. doi: 10.3389/fncel.2018.00221.

Briançon-Marjollet, A., Ghogha, A., Nawabi, H., Triki, I., Auziol, C., Fromont, S., et al. (2008). Trio mediates netrin-1-induced Rac1 activation in axon outgrowth and guidance. Mol Cell Biol 28(7), 2314–2323. doi: 10.1128/mcb.00998-07.

Chan, C.E., and Odde, D.J. (2008). Traction dynamics of filopodia on compliant substrates. Science 322, 1687–1691. doi: 10.1126/science.1163595.

Demarco, R.S., Struckhoff, E.C., and Lundquist, E.A. (2012). The Rac GTP exchange factor TIAM-1 acts with CDC-42 and the guidance receptor UNC-40/DCC in neuronal protrusion and axon guidance. PLoS Genet 8(4), e1002665. doi: 10.1371/journal.pgen.1002665.

Dent, E.W., and Gertler, F.B. (2003). Cytoskeletal dynamics and transport in growth cone motility and axon guidance. Neuron 40, 209–227. doi: 10.1016/s0896-6273(03)00633-0.

Dominici, C., Moreno-Bravo, J.A., Puiggros, S.R., Rappeneau, Q., Rama, N., Vieugue, P., et al. (2017). Floor-plate-derived netrin-1 is dispensable for commissural axon guidance. Nature 545(7654), 350–354. doi: 10.1038/nature22331.

Forscher, P., and Smith, S.J. (1988). Actions of cytochalasins on the organization of actin filaments and microtubules in a neuronal growth cone. J Cell Biol 107(4), 1505–1516. doi: 10.1083/jcb.107.4.1505.

Fothergill, T., Donahoo, A.L., Douglass, A., Zalucki, O., Yuan, J., Shu, T., et al. (2014). Netrin-DCC signaling regulates corpus callosum formation through attraction of pioneering axons and by modulating Slit2-mediated repulsion. Cereb. Cortex 24, 1138–1151.

Giannone, G., Mege, R.M., and Thoumine, O. (2009). Multi-level molecular clutches in motile cell processes. Trends Cell Biol 19(9), 475–486. doi: 10.1016/j.tcb.2009.07.001.

Gomez, T.M., and Letourneau, P.C. (2014). Actin dynamics in growth cone motility and navigation. J. Neurochem. 129, 221–234.

Gomez, T.M., and Zheng, J.Q. (2006). The molecular basis for calcium-dependent axon pathfinding. Nat. Rev. Neurosci. 7, 115–125. doi: 10.1038/nrn1844.

Hong, K., Hinck, L., Nishiyama, M., Poo, M.M., Tessier-Lavigne, M., and Stein, E. (1999). A ligand-gated association between cytoplasmic domains of UNC5 and DCC family receptors converts netrin-induced growth cone attraction to repulsion. Cell 97, 927–941.

Ishii, N., Wadsworth, W.G., Stern, B.D., Culotti, J.G., and Hedgecock, E.M. (1992). UNC-6, a laminin-related protein, guides cell and pioneer axon migrations in C. elegans. Neuron 9(5), 873–881. doi: 10.1016/0896-6273(92)90240-e.

Jurado, C., Haserick, J.R., and Lee, J. (2005). Slipping or gripping? Fluorescent speckle microscopy in fish keratocytes reveals two different mechanisms for generating a retrograde flow of actin. Mol Biol Cell 16(2), 507–518. doi: 10.1091/mbc.e04-10-0860.

Kastian, R.F., Baba, K., Kaewkascholkul, N., Sasaki, H., Watanabe, R., Toriyama, M., et al. (2023). Dephosphorylation of neural wiring protein shootin1 by PP1 phosphatase regulates netrin-1-induced axon guidance. J Biol Chem 299(5), 104687. doi: 10.1016/j.jbc.2023.104687.

Kastian, R.F., Minegishi, T., Baba, K., Saneyoshi, T., Katsuno-Kambe, H., Saranpal, S., et al. (2021). Shootin1a-mediated actin–adhesion coupling generates force to trigger structural plasticity of dendritic spines. Cell Rep. 35(7), 109130.

Katoh, K., Hammar, K., Smith, P.J., and Oldenbourg, R. (1999). Birefringence imaging directly reveals architectural dynamics of filamentous actin in living growth cones. Mol Biol Cell 10(1), 197–210. doi: 10.1091/mbc.10.1.197.

Kennedy, T.E., Serafini, T., de la Torre, J.R., and Tessier-Lavigne, M. (1994). Netrins are diffusible chemotropic factors for commissural axons in the embryonic spinal cord. Cell 78(3), 425–435. doi: 10.1016/0092-8674(94)90421-9.

Kubo, Y., Baba, K., Toriyama, M., Minegishi, T., Sugiura, T., Kozawa, S., et al. (2015). Shootin1-cortactin interaction mediates signal-force transduction for axon outgrowth. J Cell Biol 210(4), 663–676. doi: 10.1083/jcb.201505011.

Lai Wing Sun, K., Correia, J.P., and Kennedy, T.E. (2011). Netrins: versatile extracellular cues with diverse functions. Development 138(11), 2153–2169. doi: 10.1242/dev.044529.

Li, X., Saint-Cyr-Proulx, E., Aktories, K., and Lamarche-Vane, N. (2002). Rac1 and Cdc42 but not RhoA or Rho kinase activities are required for neurite outgrowth induced by the Netrin-1 receptor DCC (deleted in colorectal cancer) in N1E-115 neuroblastoma cells. J Biol Chem 277(17), 15207–15214. doi: 10.1074/jbc.M109913200.

Lowery, L.A., and Van Vactor, D. (2009). The trip of the tip: understanding the growth cone machinery. Nat Rev Mol Cell Biol 10(5), 332–343. doi: 10.1038/nrm2679.

Mai, J., Fok, L., Gao, H., Zhang, X., and Poo, M.M. (2009). Axon initiation and growth cone turning on bound protein gradients. J. Neurosci. 29, 7450–7458. doi: 10.1523/JNEUROSCI.1121-09.2009.

Medeiros, N.A., Burnette, D.T., and Forscher, P. (2006). Myosin II functions in actin-bundle turnover in neuronal growth cones. Nat. Cell Biol. 8, 215–226.

Minegishi, T., Fujikawa, R., Kastian, R.F., Sakumura, Y., and Inagaki, N. (2021). Analyses of actin dynamics, clutch coupling and traction force for growth cone advance. J Vis Exp (176). doi: 10.3791/63227.

Minegishi, T., Uesugi, Y., Kaneko, N., Yoshida, W., Sawamoto, K., and Inagaki, N. (2018). Shootin1b mediates a mechanical clutch to produce force for neuronal migration. Cell Rep 25(3), 624–639 e626. doi: 10.1016/j.celrep.2018.09.068.

Mitchison, T., and Kirschner, M. (1988). Cytoskeletal dynamics and nerve growth. Neuron 1(9), 761–772. doi: 10.1016/0896-6273(88)90124-9.

Mogilner, A., and Oster, G. (1996). Cell motility driven by actin polymerization. Biophys. J. 71, 3030–3045. doi: 10.1016/S0006-3495(96)79496-1.

Moore, S.W., Biais, N., and Sheetz, M.P. (2009). Traction on immobilized netrin-1 is sufficient to reorient axons. Science 325, 166. doi: 10.1126/science.1173851.

Moore, S.W., Zhang, X., Lynch, C.D., and Sheetz, M.P. (2012). Netrin-1 attracts axons through FAK-dependent mechanotransduction. J Neurosci 32(34), 11574–11585. doi: 10.1523/JNEUROSCI.0999-12.2012.

Serafini, T., Kennedy, T.E., Galko, M.J., Mirzayan, C., Jessell, T.M., and Tessier-Lavigne, M. (1994). The netrins define a family of axon outgrowth-promoting proteins homologous to C. elegans UNC-6. Cell 78(3), 409–424. doi: 10.1016/0092-8674(94)90420-0.

Shekarabi, M., and Kennedy, T.E. (2002). The netrin-1 receptor DCC promotes filopodia formation and cell spreading by activating Cdc42 and Rac1. Mol Cell Neurosci 19(1), 1–17. doi: 10.1006/mcne.2001.1075.

Shekarabi, M., Moore, S.W., Tritsch, N.X., Morris, S.J., Bouchard, J.F., and Kennedy, T.E. (2005). Deleted in colorectal cancer binding netrin-1 mediates cell substrate adhesion and recruits Cdc42, Rac1, Pak1, and N-WASP into an intracellular signaling complex that promotes growth cone expansion. J Neurosci 25(12), 3132–3141. doi: 10.1523/jneurosci.1920-04.2005.

Shimada, T., Toriyama, M., Uemura, K., Kamiguchi, H., Sugiura, T., Watanabe, N., et al. (2008). Shootin1 interacts with actin retrograde flow and L1-CAM to promote axon outgrowth. J Cell Biol 181, 817–829. doi: 10.1083/jcb.200712138.

Song, H., and Poo, M. (2001). The cell biology of neuronal navigation. Nat. Cell Biol. 3, E81–88.

Suter, D.M., Errante, L.D., Belotserkovsky, V., and Forscher, P. (1998). The Ig superfamily cell adhesion molecule, apCAM, mediates growth cone steering by substrate-cytoskeletal coupling. J Cell Biol 141(1), 227–240. doi: 10.1083/jcb.141.1.227.

Suter, D.M., and Forscher, P. (2000). Substrate-cytoskeletal coupling as a mechanism for the regulation of growth cone motility and guidance. J Neurobiol 44, 97–113.

Sutherland, D.J., Pujic, Z., and Goodhill, G.J. (2014). Calcium signaling in axon guidance. Trends Neurosci. 37, 424–432.

Taylor, A.M., Menon, S., and Gupton, S.L. (2015). Passive microfluidic chamber for long-term imaging of axon guidance in response to soluble gradients. Lab on a Chip 15, 2781–2789.

Toriyama, M., Kozawa, S., Sakumura, Y., and Inagaki, N. (2013). Conversion of a signal into forces for axon outgrowth through Pak1-mediated shootin1 phosphorylation. Curr Biol 23, 529–534. doi: 10.1016/j.cub.2013.02.017.

Toriyama, M., Shimada, T., Kim, K.B., Mitsuba, M., Nomura, E., Katsuta, K., et al. (2006). Shootin1: A protein involved in the organization of an asymmetric signal for neuronal polarization. J Cell Biol 175(1), 147–157. doi: 10.1083/jcb.200604160.

Varadarajan, S.G., Kong, J.H., Phan, K.D., Kao, T.J., Panaitof, S.C., Cardin, J., et al. (2017). Netrin1 produced by neural progenitors, not floor plate cells, is required for axon guidance in the spinal cord. Neuron 94, 790–799 e793. doi: 10.1016/j.neuron.2017.03.007.

Vitriol, E.A., and Zheng, J.Q. (2012). Growth cone travel in space and time: the cellular ensemble of cytoskeleton, adhesion, and membrane. Neuron 73, 1068–1081. doi: 10.1016/j.neuron.2012.03.005.

Wu, Z., Makihara, S., Yam, P.T., Teo, S., Renier, N., Balekoglu, N., et al. (2019). Long-range guidance of spinal commissural axons by netrin1 and sonic hedgehog from midline floor plate cells. Neuron 101, 635–647 e634. doi: 10.1016/j.neuron.2018.12.025.

